# Light-induced indeterminacy alters shade avoiding tomato leaf morphology

**DOI:** 10.1101/024018

**Authors:** Daniel H. Chitwood, Ravi Kumar, Aashish Ranjan, Julie M. Pelletier, Brad T. Townsley, Yasunori Ichihashi, Ciera C. Martinez, Kristina Zumstein, John J. Harada, Julin N. Maloof, Neelima R. Sinha

## Abstract

Plants sense foliar shade of competitors and alter their developmental programs through the shade avoidance response. Internode and petiole elongation, and changes in overall leaf area and leaf mass per area, are the stereotypical architectural responses to foliar shade in the shoot. However, changes in leaf shape and complexity in response to shade remain incompletely, and qualitatively, described. Using a meta-analysis of >18,000 previously published leaflet outlines, we demonstrate that shade avoidance alters leaf shape in domesticated tomato and wild relatives. The effects of shade avoidance on leaf shape are subtle with respect to individual traits, but are combinatorially strong. We then seek to describe the developmental origins of shade-induced changes in leaf shape by swapping plants between light treatments. Leaf size is light-responsive late into development, but patterning events, such as stomatal index, are irrevocably specified earlier.

Observing that shade induces increases in shoot apical meristem size, we then describe gene expression changes in early leaf primordia and the meristem using laser microdissection. We find that in leaf primordia shade avoidance is not mediated through canonical pathways described in mature organs, but rather the expression of KNOX and other indeterminacy genes, altering known developmental pathways responsible for patterning leaf shape. We also demonstrate that shade-induced changes in leaf primordium gene expression largely do not overlap with those found in successively initiated leaf primordia, providing evidence against classic hypotheses that shaded leaf morphology results from prolonged production of juvenile leaf types.

## Introduction

Not only is the shape of a single leaf highly multivariate, but the shape of leaves within and between plants is influenced by evolutionary, genetic, developmental, and environmental factors (Chitwood et al., 2012a; 2012b; 2013; 2014; Chitwood and Topp, 2015). Over a lifetime, a plant will produce numerous leaf shapes, influenced by the development of individual leaves as their blades unequally expand (allometric expansion; Hales, 1727; Remmler and Rolland-Lagan, 2012; Rolland-Logan et al., 2014) and the different types of leaf shapes a plant produces at successive nodes, a result of the temporal development of the shoot apical meristem (heteroblasty; Goebel, 1900; Ashby, 1948; Poethig, 1990; Kerstetter and Poethig, 1998; Poethig, 2010). Therefore, leaf shape in a single plant can not be reduced to a single shape, as shapes are both ephemeral, changing from one moment to the next in individual leaves, and the shapes of leaves emerging from successive nodes not necessarily constant.

When environmental conditions induce changes in leaf shape (plasticity), it is within the abovementioned developmental context that morphology must be considered (Diggle, 2002). For example, a once prevailing hypothesis was that changes in leaf shape across successive nodes were dependent on nutrition. The rationale for this premise rested on the unique (often irregular) shapes of first emerging leaves, thought to result from abortive development because of reduced photosynthetic support from any previous leaves. Similarly, many plants produce juvenile looking leaves when shaded, interpreted again as resulting from reduced photosynthate (Goebel, 1908; Allsopp, 1954). This hypothesis has recently been revisited, as sugar has been found to be a signal mediating vegetative phase change (Yang et al., 2013; Yu et al., 2013). Careful morphological studies of leaf development can separate the different effects of shade and heteroblasty refuting these ideas, at least at a morphological level in some species. In *Cucurbita*, the changes in successively emerging leaves are morphologically observable in early leaf primordia. Despite shade-induced changes in the heteroblastic progression of mature leaf morphology, leaf primordia initiated in shade resemble those initiated in sun. This suggests that, in *Cucurbita*, shade-induced morphology results from plastic responses later in leaf development after initiation, rather than through changes in heteroblasty and timing (Jones, 1995).

In addition to responses to decreased light intensity discussed above, plants can also sense changes in light quality. Phytochrome proteins, which sense decreases in the ratio of red to far-red wavelengths in light (R:FR), initiate the shade avoidance response upon detecting deflected light from competitors (Smith and Whitelam, 1997). Shade avoiding plants typically exhibit increases in internode and petiole length, reduced leaf mass per area, alterations in stomatal patterning, and shoot/root resource reallocation as an adaptive response to overgrow competitors and better intercept light (Casal, 2012). The changes in leaf shape in response to shade are more ambiguous, and can be radically different based on morphological context (such as simple versus complex leaves) and species. For example, in *Arabidopsis*, shade avoidance is typified by greater increases in petiole length relative to the blade region and inhibited blade outgrowth (Kozuka et al., 2005; Kim et al., 2005; Tsukaya et al., 2002), but in wild relatives of domesticated tomato (*Solanum* sect. *Lycopersicon*), both the petiole and rachis region expand equally and blade outgrowth is increased. These shade-avoiding responses inversely correlate with the amount of vegetation present in the native locale of an accession, implying adaptive significance in tomato (Chitwood et al., 2012b).

Here, we begin by characterizing the effects of simulated foliar shade on leaf morphology in domesticated tomato and its wild relatives through a meta-analysis of >18,000 previous published leaflets (Chitwood et al., 2012a; 2012b; 2014). We find that the effects of decreased ratios of red to far-red wavelengths of light on leaf shape are strong, but only when multiple morphometric parameters are considered across leaflets, both within leaves and across the leaf series, of individual plants. Circularity (a measure of leaflet serration in tomato) and leaf complexity are the most strongly affected individual traits during the shade avoidance response. We then seek to determine when different shade avoiding traits manifest during leaf development by swapping plants between light treatments during early leaf development. Leaves will plastically increase their blade area late into development if moved into simulated foliar shade from an initial sun light treatment, but other traits, such as stomatal patterning, are irrevocably specified earlier during development. Observing increases in shoot apical meristem size under low red to far-red light conditions, we then perform laser-capture microdissection (LMD) to analyze effects of simulated foliar shade on gene expression in the first emerging leaf primordium (P1) and the meristem. The most conspicuous change in gene expression is increased expression of *LeT6* (the tomato ortholog of *SHOOTMERISTEMLESS*) and other indeterminacy-related genes in the P1, consistent with increases in leaflet serration and leaf complexity observed in tomato during the shade avoidance response. Finally, to determine the developmental context of our observations, we compare gene expression changes during shade avoidance with heteroblastic gene expression (i.e., successively initiating leaves at the same developmental stage) in the shoot apical meristem and young leaf primordia. Gene expression induced by decreased red to far-red ratios in light and that which changes with progression through the heteroblastic series in leaf primordia are largely distinct, suggesting not only that shade avoidance response is not mediated through heteroblastic changes, but also that increase in leaf complexity in these two contexts employs distinct suites of genes.

## Results

### Morphology of shade avoiding tomato leaves

Increases in internode and petiole length induced during the shade avoidance response are well known and occur in tomato (Fig. 1A). With regard to leaf shape, we previously described increases in blade area, and elongation throughout all parts of the leaf, petiole and rachis alike, in shade avoiding tomato plants (Chitwood et al., 2012a). When more comprehensive measures of leaflet contours are assessed using Elliptical Fourier Descriptors (EFDs; a method to globally describe closed contours as a Fourier series, see Kuhl and Giardina, 1982 and Iwata et al., 2002), across successively emerging leaves and the leaf proximal-distal axis, shade avoidance does not induce “shade morphology” per se. Rather, the developmental trajectory of leaf shapes throughout the leaf series takes on qualities associated with more adult leaves with accelerated development (Chitwood et al., 2012a). These shape changes are subtle, and may ultimately reflect small accelerations in the transition to reproduction. Shape changes induced by shade are so subtle that when we assessed leaf shape in a set of 76 near-isogenic introgression lines harboring tiled genomic segments from the desert adapted tomato relative *Solanum pennellii* (referred to hereafter as the “ILs”), shade effects for univariate traits in individual leaflets were not significant and were subsequently ignored in models (Chitwood et al., 2014).

**Figure 1:**
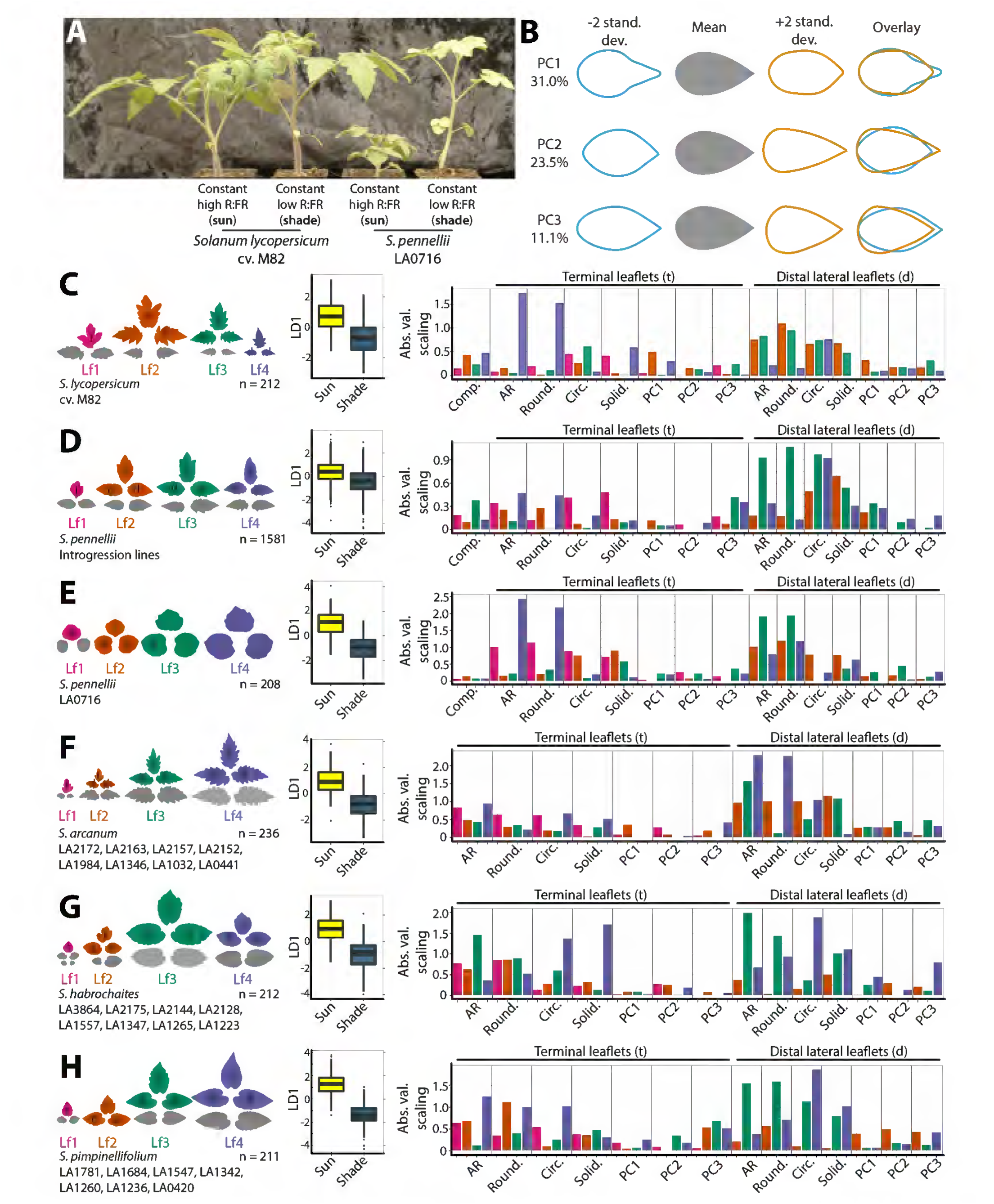
The morphology of shade avoiding leaves in domesticated tomato and wild relatives. **A)** *Solanum lycopersicum* cv. M82 (domesticated tomato) and *S. pennellii* LA0716 grown under high (simulated sun) and low (simulated foliar shade) red to far-red light conditions. *S. pennellii* has an exaggerated shade avoidance response compared to domesticated tomato, characterized by increased internode and petiole lengths. **B)** Eigenleaves representing theoretical leaf shapes found-/+ 2 standard deviations along each principal component. Percent variance explained by each principal component indicated. Principal component space and other morphometric parameters calculated from previously published data on wild relatives of tomato (Chitwood 2012a; 2012b) and field-(Chitwood et al., 2013) and chamber-grown (Chitwood et al., 2014) *S. pennellii* introgression lines and parents. Eigenleaves reproduced here from Chitwood et al., 2014 for meta-analysis purposes. **C-H)** Separate Linear Discriminant Analyses (LDAs) by light treatment performed for leaflets across the leaf series for **C)** *S. lycopersicum* cv. M82, **D)** *S. pennellii* ILs, **E)** *S. pennellii* LA0716, **F)** *S. arcanum*, **G)** *S. habrochaites*, and **H)** *S. pimpinellifolium* accessions. Leaflets used in analyses are indicated by color (magenta = leaf 1; orange = leaf 2; teal = leaf 3; lavender = leaf 4) and text (“t” = terminal; “d” = distal lateral). Boxplot shows resulting LD1 values for each analyses by light treatment (yellow = simulated sun; dark blue = simulated foliar shade). Bar plots show the absolute values of scalings resulting from LDAs, which indicate the relative contributions of traits to the discrimination of leaflets by light treatment.

Here, before embarking upon describing the developmental gene regulatory networks in the shoot apical meristem and early leaf primordia that are activated during shade avoidance in domesticated tomato, we wish to fully describe the morphological changes in tomato leaf shape in a meta-analysis. We draw upon 1) >11,000 sampled leaflets from up to eight accessions each of *Solanum arcanum*, *S. habrochaites*, and *S. pimpinellifolium* (Chitwood et al., 2012a; 2012b) and 2) >33,000 leaflets from the *S. pennellii* ILs and their two parents, *S. lycopersicum* cv. M82 and *S. pennellii* LA0716 (Chitwood et al., 2014; **Dataset S1**). Leaflets from both datasets were grown under simulated sun and foliar shade conditions with equal PAR but lower red-to-far red wavelength ratios in shade conditions created by alternating white light and far-red fluorescent bulbs (see Materials and Methods). A Principal Component Analysis (PCA) describing EFDs of these leaflets and field-grown leaves from the ILs (Chitwood et al., 2013) was previously calculated to describe the overall tomato leaf morphospace (Chitwood et al., 2014), and we use the previously published PCs here in our analysis (Fig. 1B). In addition to PCs, shape is described by measures influenced by length-to-width ratio, including aspect ratio (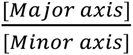 of the best-fitted ellipse) and roundness(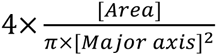, inversely related to aspect ratio), and traits that largely reflect serrations and lobing, including circularity(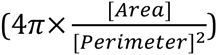) and solidity(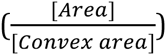).

Because shade effects on leaf morphology are so subtle for individual shape features in each measured leaflet (Chitwood et al., 2014), we have used a Linear Discriminant Analysis (LDA) to maximize the discrimination of leaf shapes from simulated sun and foliar shade conditions. We began by considering each trait alone across 7 different leaflet types (the terminal leaflets of the first 4 leaves and averages of distal lateral leaflets from leaves 2-4; >18,000 leaflets). Because of extremely high replication within single genotypes or classes of genotypes, the resulting linear discriminant (LD1) is significantly different between sun and shade treatments for nearly all traits in all genotype classes (Table 1). However, the ability to discriminate shade and sun leaves is not nearly as powerful for single traits when all traits are considered in combination across the leaf series (Table 1). Previous analyses suggested that the morphological effects of shade avoidance on tomato leaves were either subtle (Chitwood et al., 2012a) or not significant (Chitwood et al., 2014) when individual traits were considered alone for particular leaflets. However, using the resultant linear discriminant space defined for all measured traits across the leaf series in a predictive fashion, a majority of plants, across tomato and its wild relatives, can be correctly identified as growing under simulated sun or foliar shade conditions (Table 2). These results suggest that although morphological changes induced by shade for any single shape feature for a particular leaflet are weak, that the signature across the leaf series and over many shape traits is combinatorially quite strong.

**Table 1:**
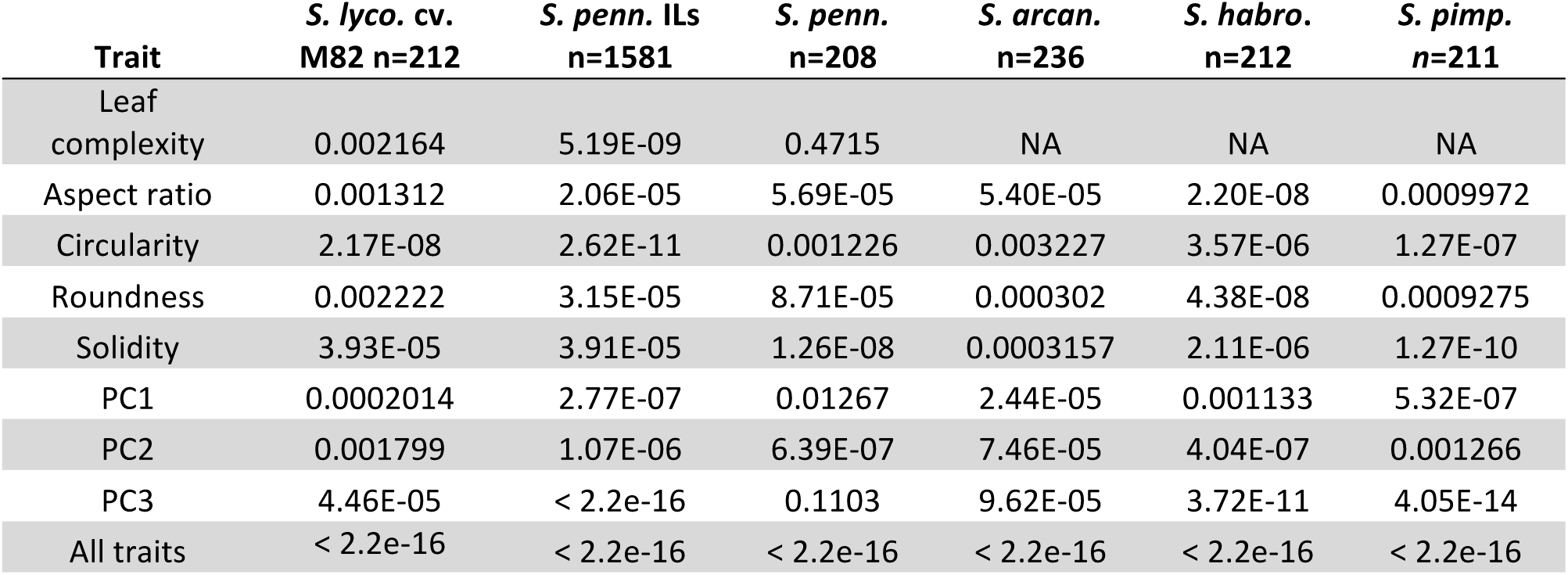
p-values for differences in LD1 values between light treatments. LDAs were performed for the given trait for leaflets across the leaf series for the indicated genotypes. Shown are p-values for differences in LD1 values between simulated sun and foliar shade light treatments as calculated using the Mann-Whitney U test. Results from an LDA performed using all traits is also provided.

**Table 2:** Predicted identities of leaflets. Using calculated Linear Discriminants, plants were predicted as originating from simulated sun or foliar shade treatments using morphometric trait information for leaflets across the leaf series. Shown are numbers for the actual and predicted light treatments for plants, and the percent correct identification for each light treatment.

| Species | Number of plants | Shade predicted as shade | Sun predicted as sun | Shade predicted as sun | Sun predicted as shade | Correctly predicted shade | Correctly predicted sun |
| --- | --- | --- | --- | --- | --- | --- | --- |
| *S. lyco. cv. M82* | 212 | 82 | 80 | 25 | 25 | 76.6% | 76.2% |
| *S. penn. ILs* | 1581 | 506 | 548 | 278 | 249 | 64.5% | 68.8% |
| *S. penn.* | 208 | 97 | 89 | 8 | 14 | 92.4% | 86.4% |
| *S. arcan.* | 236 | 101 | 87 | 23 | 25 | 81.4% | 77.7% |
| *S. habro.* | 212 | 86 | 89 | 18 | 19 | 82.7% | 82.4% |
| *S. pimp.* | 211 | 87 | 103 | 13 | 8 | 87.0% | 92.8% |

What are the discriminants that separate sun and shade avoiding leaflets? An analysis of the absolute values of the LD1 scalings (coefficients that when multiplied by trait values additively produce the resultant linear discriminant values) for each genotypic class (Fig. 1C-H) shows that shape attributes describing length-to-width ratio (aspect ratio and roundness) and serration and lobing (circularity and solidity), but to a lesser extent PCs derived from EFDs, all contribute to the discrimination of shade and sun plants (**Dataset S2**). These contributions vary across the leaf series, sometimes strongly, for different genotype classes. From these results, to say that a particular aspect of shape, or particular leaflets in the leaf series, predominately contribute to shade avoiding leaf morphology would be premature. Rather, shade avoidance strongly affects leaf shape in a way that touches upon many metrics commonly used to quantify morphology.

To demonstrate the subtle contributions of individual morphometric parameters to shade avoiding leaf morphology leaf complexity, aspect ratio, roundness, circularity, and solidity were modeled for *S. lycopersicum* cv. M82 data as a function of leaflet type, light treatment, and their interaction using Analysis of Variance (ANOVA). All morphometric traits strongly vary across leaflet types (or the leaf series, for leaf complexity), but only leaf complexity and circularity (a measure of serration in tomato leaflets) significantly vary by light treatment in domesticated tomato, as defined by an alpha level of p < 0.05 (Table 3). Additionally, the interaction term between leaflet type and light treatment is significant for circularity as well. Boxplots for leaf complexity and circularity demonstrate the subtlety of the effects of light treatment on domesticated leaf morphology when considered within the greater context of shape changes induced by the leaf series itself (Fig. 2). The directions of these effects are such that shade avoidance increases leaf complexity (Fig. 2A) and increases leaflet serration (lower circularity values), especially in distal lateral leaflets compared to terminal leaflets (Fig. 2B). We first describe the morphogenetic window during which the shade avoidance response manifests developmentally in leaves and then describe gene expression changes in the shoot apical meristem and early leaf primordia induced by shade consistent with these morphological effects.

**Table 3:** Light affects domesticated tomato leaf complexity and serration. For each of the indicated traits, an ANOVA was performed considering leaflet type (or leaf, for leaf complexity), light treatment, and a leaflet:treatment interaction effect. Only significant terms were included in the final model, for which p-values are indicated. NS, not significant at 0.05 alpha level.

| Trait | Leaflet | Light treatment | Interaction |
| --- | --- | --- | --- |
| Leaf complexity | < 2.2E-16 | 0.000103 | NS |
| Aspect ratio | < 2.2E-16 | NS | NS |
| Roundness | < 2.2E-16 | NS | NS |
| Circularity | 2.17E-13 | 1.09E-02 | 0.0379 |
| Solidity | < 2.2E-16 | NS | NS |

**Figure 2:**
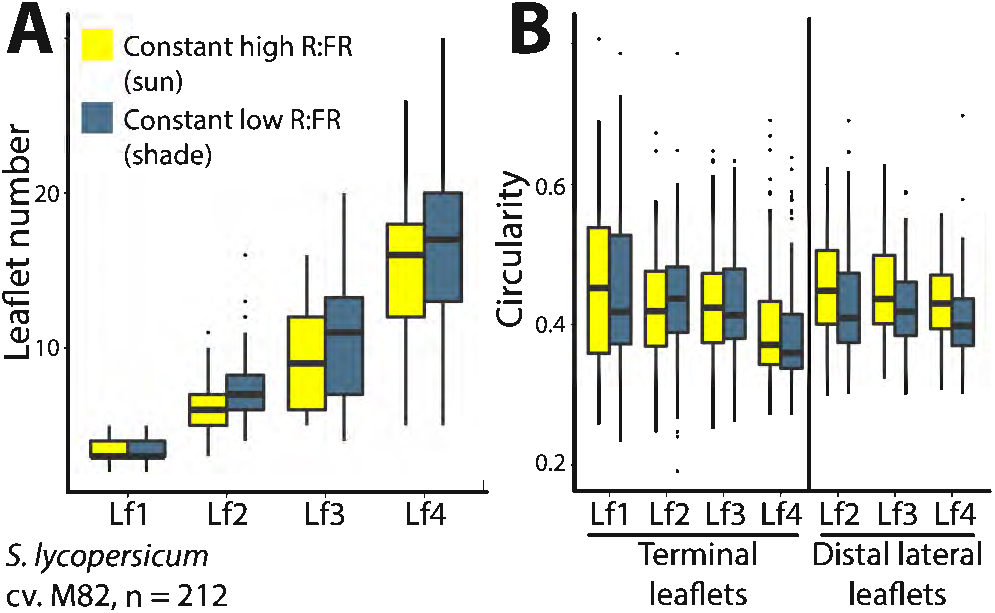
Leaf complexity and serration are increased during the shade avoidance response in domesticated tomato. Boxplots showing **A)** leaflet number, a measure of leaf complexity, and **B)** circularity, decreased values of which indicate increased serration in tomato leaflets, across leaves and leaflet types in domesticated tomato. Not only does shade avoidance increase leaf complexity and serration, but also the heteroblastic series (developmental differences in successively emerging leaves). See Table 3 for significance values of light and developmental effects. Light treatment indicated by color (yellow = simulated sun; dark blue = simulated foliar shade).

### Early vs. late developmental responses to shade

To resolve the developmental window during which shade avoidance traits manifest in tomato leaves, plants were reciprocally swapped between simulated sun (high R:FR) and foliar shade (low R:FR) conditions at 20 dap (days after planting). At 20 dap, leaf 2 has initiated but has yet to fully expand in plants grown under either constant sun or shade conditions (Fig. 3A). Comparing plants grown under constant sun (yellow) and shade (dark blue) conditions reveals shade-induced phenotypes. If plants swapped from sun into shade (orange) or from shade into sun (cyan) retain the phenotype associated with their initial growth conditions, it suggests that shade alters the trait during early leaf development and that it persists regardless of subsequent changes in light quality. Traits that are altered by the new environment post-swap indicate that a trait can be changed late in development (after laminar expansion). As described below, examples of shade-induced traits determined pre-swap include stomatal patterning (stomatal index, the ratio of stomata to other epidermal cells, distinct from stomatal density) and the size of the shoot apical meristem, which are irrevocably altered by pre-swap light conditions (Fig. 3B).

**Figure 3:**
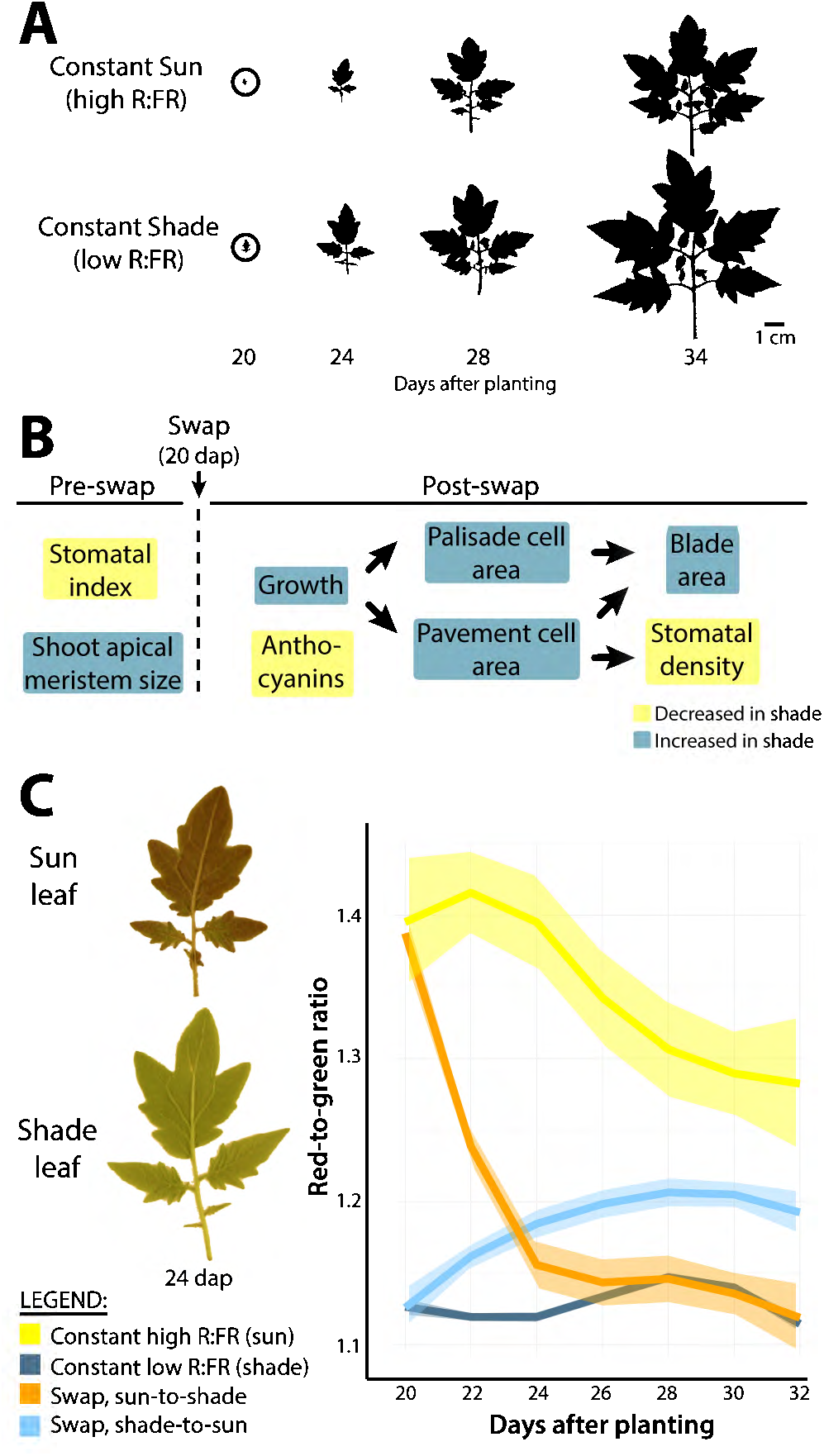
Swap experiments resolve shade avoidance responses before and after laminar expansion. **A)** Outlines of leaf 2, in constant simulated sun (high R:FR) and constant simulated shade (low R:FR) light treatments, show the exponential increases in growth that occur between 20-34 days after planting (dap) during which traits were recorded. In swapped light conditions, plants are swapped into different light treatments at 20 dap, at the onset of laminar expansion in leaf 2. Traits specified pre-and post-swap are indicated. Arrows indicate the developmental progression of traits, that is the effects of one trait (e.g., cell expansion) on subsequent traits (e.g., blade area and stomatal density). Yellow indicates decreased traits value upon swap to shade, dark blue increased values. **C)** Red-to-green ratio in leaves is influenced by shade 20 dap. Sun leaves swapped into shade can dissipate anthocyanins such that levels comparable to constant shade plants are attained. Transparent bands indicate 95% confidence interval of loess regression. Yellow = constant sun; dark blue = constant shade; orange = sun-to-shade; cyan = shade-to-sun.

Post-swap, shade-induced increases in leaf size are achieved through increased palisade and pavement cell size, which increase blade area and reduce absolute stomatal density (Fig. 3B, arrows indicate the developmental progression of post-swap shade-induced phenotypic changes; e.g., cellular expansion leads to increased blade area). The ability of a post-initiated leaf from one light condition to achieve the attributes of leaves from the other condition indicates the trait can be influenced late in development, after blade expansion. An excellent example of this scenario is anthocyanin accumulation under simulated sun conditions, which dissipates upon shifting a mature leaf into shade (Fig. 3C).

Overall blade area in leaves is remarkably labile late into development. Leaves swapped from sun into shade (orange) significantly increase in overall blade area compared to leaves that remain in sun (yellow), and vice versa (Fig. 4A).

**Figure 4:**
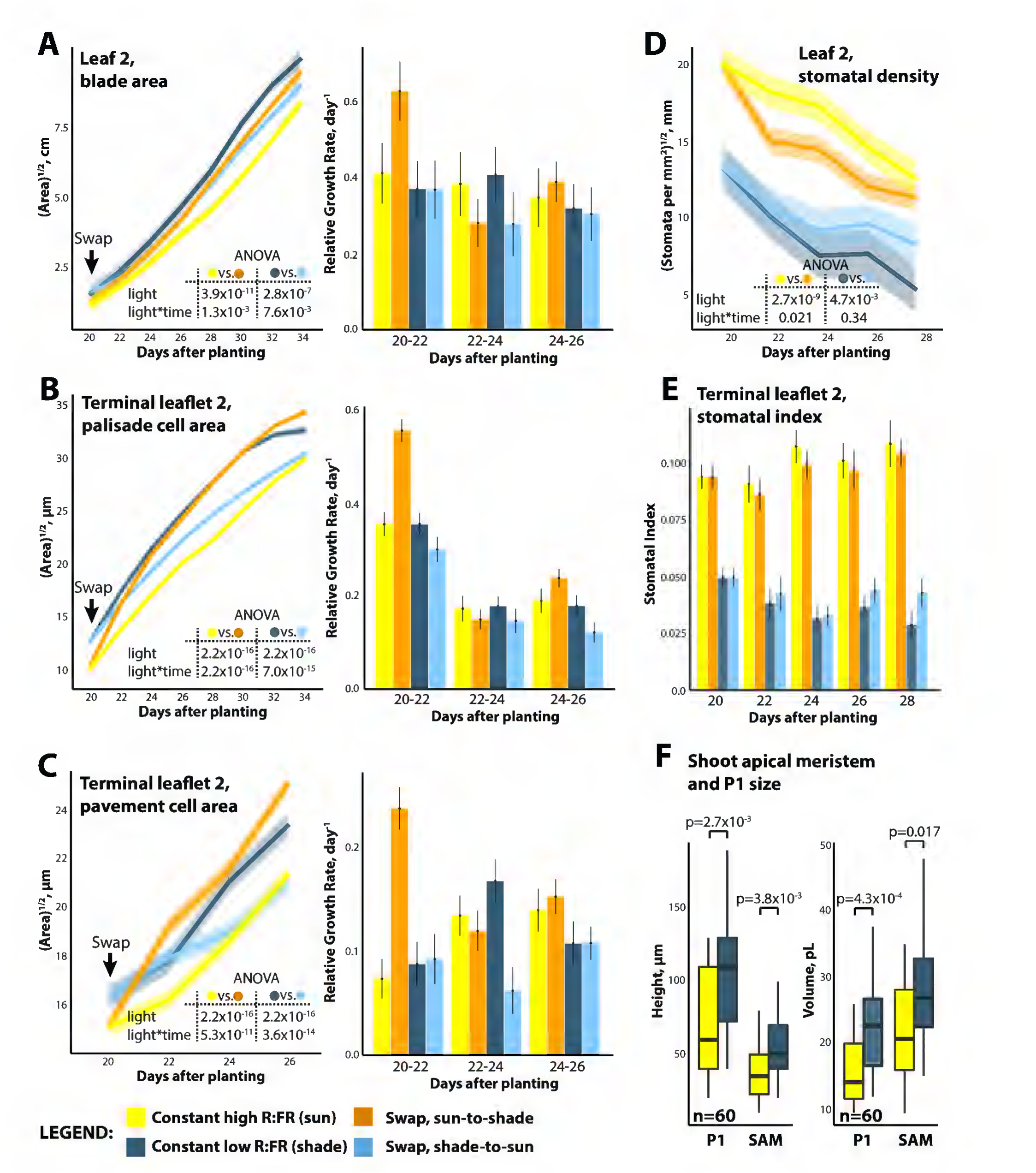
Shade avoidance is characterized by rapid increases in relative growth rate upon acute shade exposure. **A-C)** Swap results for leaf 2 **A)** overall blade area, **B)** palisade cell area, and **C)** pavement cell area. Indicated for each trait are p values for factors from fitted ANOVA models comparing constant sun to sun-to-shade and constant shade to shade-to-sun treatments. For each trait, relative growth rates for each condition over 2 day intervals are shown. **D-E)** Swap results for leaf 2 stomatal traits. **D)** Absolute stomatal density and **E)** stomatal index (patterning) from the terminal leaflet. **F)** Increases in P1 and SAM height and volume as measured from reconstructed histological sections. p values indicate significant differences between constant sun and shade treatments for each trait measured. Yellow = constant sun; dark blue = constant shade; orange = sun-to-shade; cyan = shade-to-sun.

Importantly, the blade area of swapped leaves does not fully assume the area of leaves grown under constant conditions, suggesting that although labile, a fraction of the size of leaf 2 is fixed before 20 days, when blade expansion had yet to occur. Like blade area, cell sizes (both palisade and pavement cells) in swapped leaves rapidly adjust to the post-swap light condition, but unlike blade area, attain the size of cells in leaves constantly grown under the post-swap light condition (Fig. 4B-C).

Calculation of relative growth rate (the change in trait value over time normalized for the magnitude of the trait) shows that growth under all conditions occurred at similar relative rates except for a “burst” of growth when plants are first swapped from sun into shade (Fig. 4A-C). That is, despite plants grown under constant shade conditions being larger, they still grow at equivalent relative growth rates to sun plants once size is taken into account. The “burst” in relative growth rate upon first exposure to foliar shade is rapid, occurring within the first two days after the swap, and transient, subsiding soon after. The rapid expansion of cells introduced to shade, in fully patterned leaves in which most cell division has subsided, contributes towards the increases in blade area observed in post-swap shade-avoiding plants. Further experiments will determine for how long during development a leaf can respond to shade subsequently after laminar expansion, but the response may ultimately be associated with the initial exponential growth rate of leaves and begin to subside as leaf growth plateaus and ultimately ceases.

Some leaf traits are fixed before the 20 day swap and irrevocably changed despite any post-swap changes in light quality encountered. Stomatal density (the number of stomata per given leaf surface area) is higher in sun compared to shade conditions, and inversely tracks pavement cell size in swapped plants, suggesting that shade-induced increases in pavement cell area “push” stomata away from each other after the swap (Fig. 4D). However, stomatal index (the ratio of stomata to all epidermal cell types, a measure of stomatal patterning), which is also higher in sun compared to shade, remains constant after swapping, indicating that the ratio of stomata to total epidermal cell number had been previously, and irrevocably, specified previous to the swap during early leaf development (Fig. 4E).

Previous experiments, purposefully altering either CO_2_ concentrations (Lake et al., 2001) or irradiance (Thomas et al., 2004) of mature leaves, have observed non-cell autonomous effects on stomatal patterning, epidermal cell morphology, and overall size of developing leaves. Such an observation is consistent with mature leaves sensing changes in light quality and transmitting signals (whether metabolites, hormones, transcripts, or proteins, or a combination thereof) to alter development in young leaf primordia, similar in concept to the pre-swap changes in leaf morphology we observe (Figs. 3-4). Shade avoidance may require a similar mechanism: although the shoot apical meristem of tomato is relatively exposed compared to other species, it is thoroughly encapsulated by leaf primordia, preventing the direct reception of light possible in mature leaves. An important first step towards understanding the connection between shade perception and developmental change is to ask whether leaf primordia in the shoot apical meristem are altered during the shade avoidance response.

Analysis of the volume and height of the P1 (plastochron 1, designating the first separated, emerging leaf primordium from the meristem) and SAM (shoot apical meristem, in this case referring to the meristem and P0, or the incipient leaf) shows both are significantly larger in continuous shade compared to sun (Fig. 4F). As the shoot apical meristem includes the incipient leaf, this observation shows that, in addition to shade-induced changes we observe later during development, shade likely alters morphology from the very beginnings of leaf development as well.

### Shade alters developmental programs in the shoot apical meristem

To determine the gene expression changes underlying 1) the global changes in leaf morphology we observe (Fig. 1), especially those increasing leaflet serration and leaf complexity (Fig. 2; Table 3), and 2) increases in leaf primordium and meristem size induced by shade (Fig. 4F), we captured the shoot apical meristem (SAM, which contains both meristem proper and the incipient leaf) and the first separated leaf primordium (P1) using laser microdissection (Fig. 5A). Given the rapid and temporary “burst” of growth observed in relative growth rate (Fig. 4A-C), we measured gene expression in the meristem and organ primordia from plants grown in constant high R:FR (sun) and counterparts shifted from high R:FR to low R:FR light (shade) for 28 hours, assuming that the largest changes in gene expression would correspond to the acute changes in relative growth rate we observe (Fig. 4).

**Figure 5:**
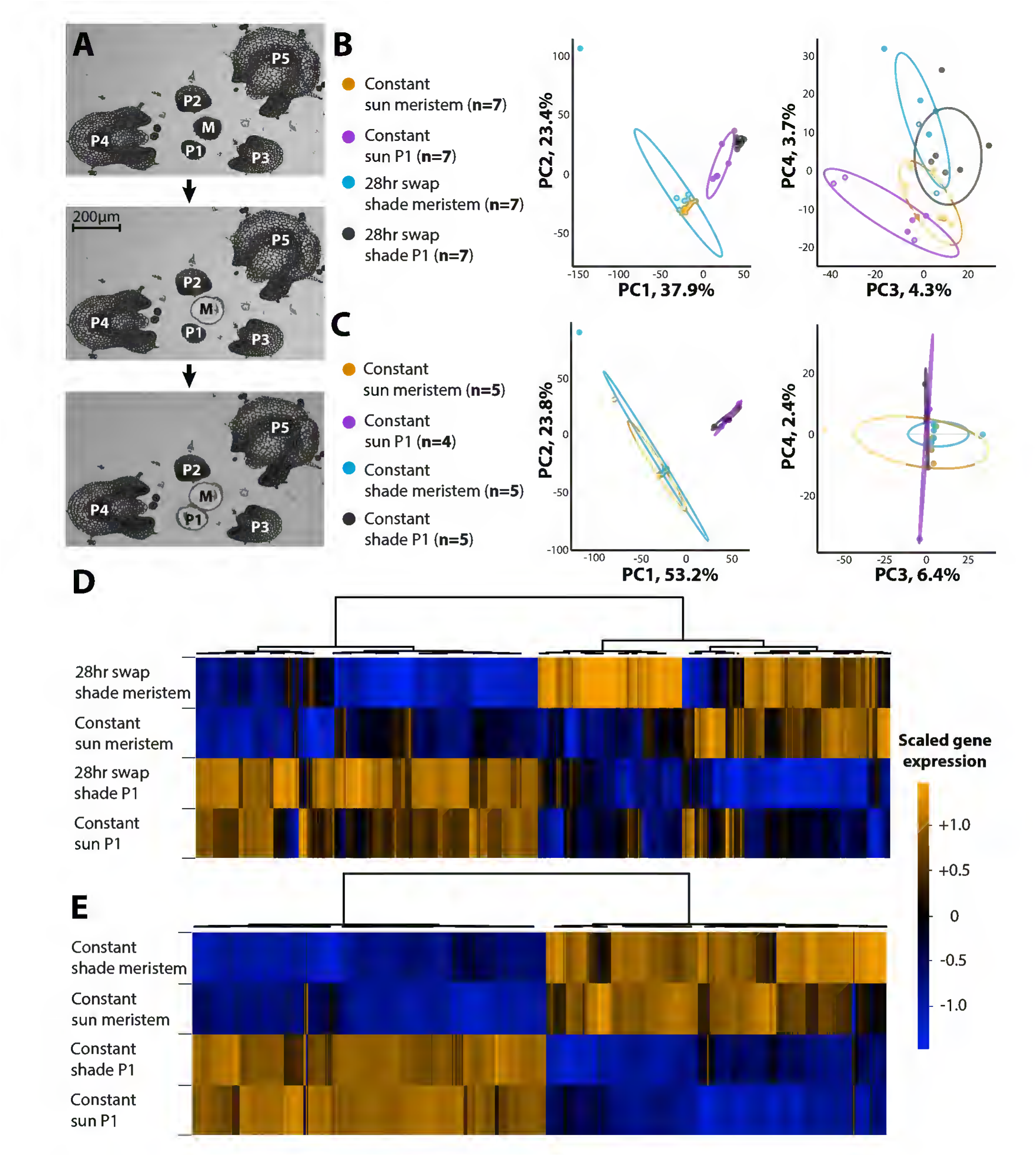
Transcriptional responses to shade in the shoot apical meristem. **A)** Steps of laser microdissection, capturing the SAM (denoted “M” for meristem”) followed by P1. **B-C)** Principal component analysis (PCA) on replicate gene expression values. **B)** Constant sun and 28 hr swap to shade and **C)** constant sun and constant shade conditions. Note the separation by light conditions in PCs 3 and 4 under the swap experiment absent under constant conditions. Ovals indicate 95% confidence ellipses. Orange = constant sun SAM; magenta = constant sun P1; cyan = constant/28 hr shade SAM; black = constant/28 hr shade P1. **D-E)** Hierarchical clustering of those genes differentially expressed either between tissues or by light treatments. **D)** Constant sun and 28 hr swap to shade and **E)** constant sun and constant shade conditions. Note that under constant light conditions, most differential expression separates between meristem and primordium, whereas in the swap experiment light conditions contribute to gene expression differences as well. Color indicates averaged, scaled gene expression levels across tissues and treatments. Orange, high expression; blue, low expression.

Using a Principal Component Analysis (PCA), PC1 and PC2 cluster RNA-Seq replicates more strongly by organ type (SAM or P1) rather than light treatment (constant sun or 28 hours shade), indicating the differentiation of a leaf from the meristem alters gene expression more strongly than light conditions (Fig. 5B).

Effects of light treatment are still observable, however, as PC3 and PC4 cluster samples more closely by light treatment than organ type.

Acute changes in shoot apical meristem (SAM) gene expression upon exposure to 28 hours shade relative to constant sun grown controls are modest. Only 54 genes are differentially expressed in the SAM, and a majority of those (52) are down regulated upon the shift to low R:FR (**Dataset S3**). Among these genes are *LIGHT-SENSITIVE HYPOCOTYL* (*LSH*) and *TORNADO* (*TRN*) homologs (as identified through reciprocal BLAST searches with *Arabidopsis* genes, described in Chitwood et al., 2013). *LSH* genes mediate light-regulated development, meristem maintenance, and organogenesis (through boundary specification; Zhao et al., 2004; Cho et al., 2011). In particular, in tomato the *LSH6* homolog *TERMINATING FLOWER* leads to premature meristem growth arrest and the *LSH3b* homolog is implicated in modulating leaf complexity across the tomato clade (MacAlister et al., 2012; Ichihashi et al., 2014), although neither of these genes is the *LSH* homolog we observe differentially expressed. Both tomato *LSH6* and *LSH3b* homologs interact with BOPa, which patterns primary leaflet specification across the tomato rachis, and *LSH3b* directly binds the promoter of *PETROSELINUM* that regulates the tomato ortholog of *SHOOTMERISTEMLESS* (*LeT6*) which modulates leaf complexity (Janssen et al., 1998; Kimura et al., 2008; Ichihashi et al., 2014). *trn* mutations disrupt auxin regulation and promote senescence (Cnops et al., 2006). *TRN* function is epistatic to *PHANTASTICA*/*ASYMMETRIC LEAVES1* (*PHAN*/*AS1*), which down-regulates KNOX (knotted-1 like homeobox) gene expression in the incipient leaf. Thus, meristems acutely exposed to low R:FR light conditions (shade) for 28 hours exhibit changes in the expression of genes intimately associated with leaf development and complexity in tomato, which are altered by the shade avoidance response (Figs. 1-2; Table 3).

The P1 response to acute shade is far more robust than that of the SAM. Genes are both significantly down and up regulated (290 down, 355 up) in the P1 (**Dataset S4**). In the P1, acute shade promotes the expression of genes associated with indeterminacy, cell proliferation, and the meristem while down-regulating genes associated with leaf differentiation and patterning.

Up regulated genes in the P1 upon acute shade exposure are significantly enriched for an auxin GO (gene ontology) term and numerous transcription-related terms (**Dataset S5**). Many of the genes described by transcriptional GO terms are developmental regulators associated with meristem identity and, in tomato, leaf complexity. Notably, of the 10 most differentially expressed genes in the shade P1, five are KNOX and KNOX-related, including *LeT6* (*Lycopersicon esculentum T6*; **Dataset S4**). Also highly expressed in the shade P1 are *WUSCHEL* (*WUS*), *CLAVATA1* (*CLV1*), *HANABA TANARU*, and *PLETHORA* (*PLT*) and *EPIDERMAL PATTERNING FACTOR-LIKE* (*EPFL*) homologs, the former consistent with increases in leaf indeterminacy and complexity (Fig. 2; Table 3) and the latter consistent with shade-induced changes in stomatal patterning (Fig. 4D-E). Commensurately, genes promoting leaf differentiation are down regulated, including *PHAN*/*AS1*, *SAWTOOTH (SAW)*, *JAGGED*/*LYRATE* (*JAG*/*LYR*), and TCPs (TEOSINTE BRANCHED1, cycloidea and PCF transcription factors) (**Dataset S4**).

Our results demonstrate that key regulators of meristem identity specifying indeterminacy are highly up regulated, while complimentary promoters of leaf differentiation and patterning down regulated, in the P1 upon shifting to shade for 28 hours. The up-regulation of KNOX gene expression early in leaf development (at the P0-P1 stage) is important. In tomato (and many other species), KNOX gene activity promotes leaflet formation (Janssen et al., 1998; Bharathan et al., 2002).

Additionally, KNOX activity is associated with meristem identity (Vollbrecht et al., 1991) and in leaves has stage specific effects. In young tomato leaf primordia, KNOX activity prolongs the process of leaf initiation, consistent with the changes in gene expression we observe associated with an indeterminate fate and antagonistic with differentiation (Shani et al., 2009). Again, such changes in gene expression are consistent with increases in leaflet serration and leaf complexity we observe induced during the shade avoidance response (Figs. 1-2; Table 3).

With respect to hormones, we observe increased expression of *AUXIN RESISTANT* (*AUX*), IAAs (INDOLE-3-ACETIC ACID INDUCIBLEs), and ARFs (AUXIN RESPONSE FACTORs), as well as increased cytokinin oxidase (*CKX*) homolog transcript levels in the P1 samples exposed to acute shade (**Dataset S4**). Auxin-induced cytokinin breakdown has been shown to lead to a transient arrest in leaf development in *Arabidopsis* (Carabelli et al., 2007). However, unlike the inhibited blade outgrowth observed in *Arabidopsis*, shade induces increased laminar outgrowth in tomato (Chitwood et al., 2012b), suggesting alternative regulation of cytokinin during shade avoidance in tomato. We observe increased cytokinin responsiveness through increased expression of cytokinin-inducible *WOODEN LEG*, and numerous *HISTIDINE-CONTAINING PHOSPHOTRANSMITTER* (*AHP*) and ARR (Arabidopsis response regulator) homolog transcripts and cell division markers, consistent with the increased expression in KNOX and other meristem identity genes described above. KNOX gene activity and cytokinin are known to affect SAM size (Vollbrecht et al., 2000; Guiliani et al., 2004; Leibfried et al., 2005), providing a basis for the shade-induced increases in SAM and P1 volumes we observe (Fig. 4F).

To better understand the gene regulatory networks that underlie acute shade-induced increases in SAM cell proliferation, growth, and indeterminacy, promoters of differentially expressed genes were analyzed for over-represented motifs. Many promoters of differentially expressed genes contain multiple types of significantly enriched motifs (Fig. 6A). BELLRINGER (a Class I KNOX gene transcriptional regulator) and ARF binding sites are among enriched motifs, congruent with KNOX and auxin response genes that are differentially expressed upon exposure to acute shade in the P1. Interestingly, Evening element and CCA1 (CIRCADIAN CLOCK ASSOCIATED1) bindings sites are overrepresented, which reflects the gating of shade avoidance response by circadian rhythms (Salter et al., 2003; Nozue et al., 2011). We also observe an enrichment of abscisic acid (ABA) motifs, reflecting the increased of expression of ABA-related transcripts (e.g., *LEA* (*LATE EMBRYOGENESIS ABUNDANT*) and *AREB* (*ABA-RESPONSIVE ELEMENT BINDING PROTEIN*) homologs) in the shade P1 and increased ABA levels in shaded tomato leaves (Cagnola et al., 2012).

**Figure 6:**
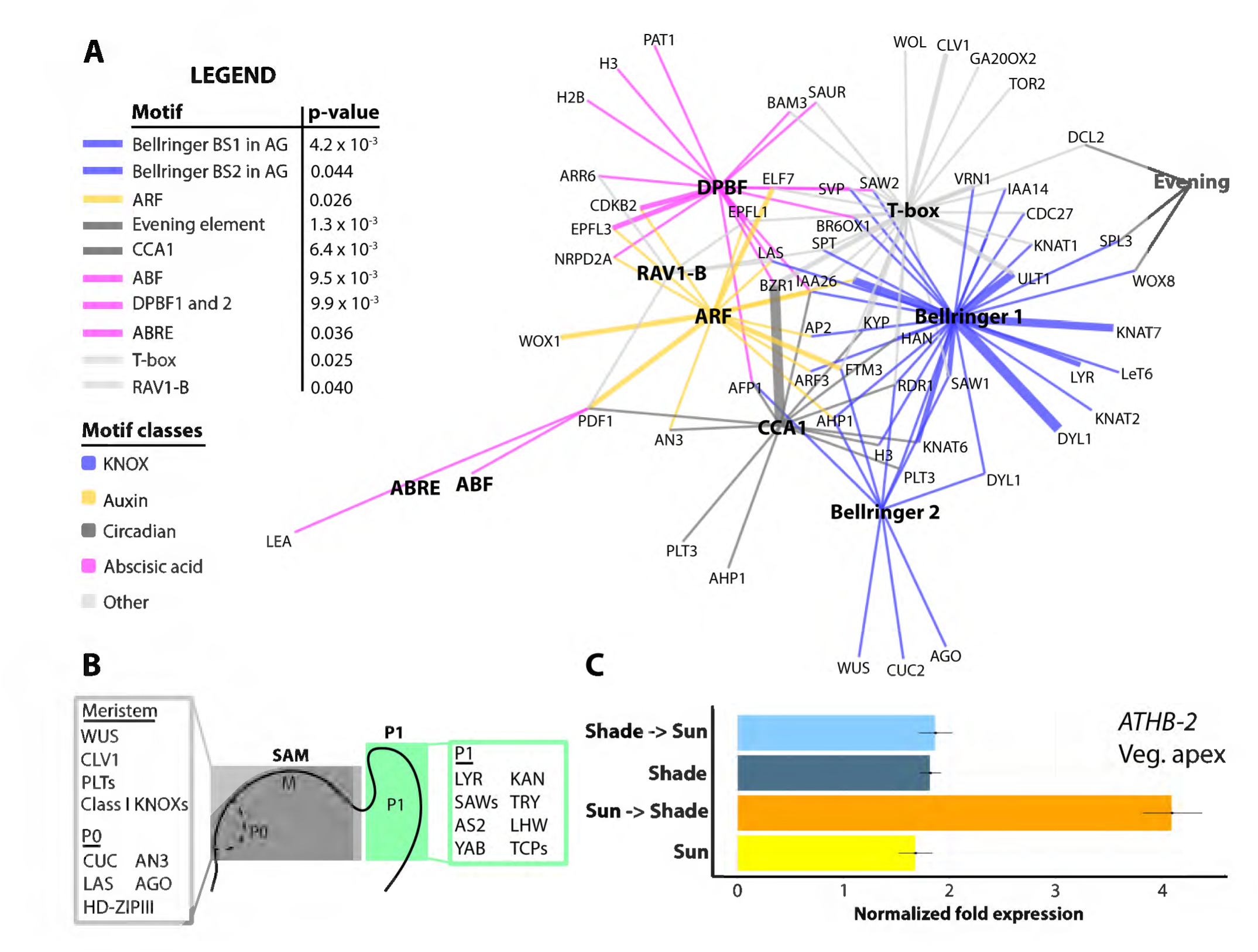
Enriched motifs in promoters of select genes acutely up regulated in the shade P1 and *ATHB-2* expression. **A)** Provided are p values for enrichment of different motifs, color indicating different motif classes. Network shows select developmental regulators (terminal nodes) that harbor enriched motifs (hubs, bold). Promoters of many developmental regulators contain motifs of many classes. Classes of motifs: blue = KNOX, yellow = auxin, dark gray = circadian, magenta = abscisic acid, light grey = other. Edge thickness is proportional to number of motifs present in the promoters of the respective genes. **B)** Select genes with significantly increased expression in respective tissues, regardless of light treatment, are shown (SAM, grey; P1, green). As expected, numerous regulators of indeterminacy are up regulated in the SAM, whereas transcripts promoting leaf differentiation and patterning are expressed at higher levels in the P1. Many of the genes associated with indeterminacy in the SAM are differentially expressed upon acute shade treatment in the P1 (see Fig. 7E). **C)** *ATHB-2* is rapidly, and transiently, up regulated after exposure to low R:FR light treatment for 28 hours in vegetative apices. This differential expression is absent in LCM SAM and P1 samples. Vegetative apices contain P5-7 leaf primordia, that are mm in length, demonstrating that differential expression is restricted to relatively mature leaves. Error bars indicate standard deviation. Yellow = constant sun; dark blue = constant shade; orange = sun-to-shade; cyan = shade-to-sun.

### Shade avoidance in the shoot apical meristem is ephemeral

Our own results (Fig. 4A-C) and previous work demonstrates that the shade avoidance response is a rapid and transient phenomenon (Sessa et al., 2005). If instead of microdissecting SAM and P1 tissue shifted to shade for only 28 hours, samples are collected from constant sun and shade conditions (Fig. 5C), transcriptional responses to shade are largely abolished. The absence of a prominent transcriptional shade avoidance response under constant light conditions reveals that, molecularly, shade avoidance in the tomato SAM and P1 is ephemeral. The differences in shade avoidance response under transient versus constant shade conditions are easily visualized by hierarchical clustering (Fig. 5D-E). When plants are exposed to only 28 hours of shade, expression patterns of differentially expressed genes separate by tissue and light treatment (Fig. 5D), whereas under constant light conditions, expression patterns are mainly explained by tissue type alone (Fig. 5E). No significant differential gene expression between constant high and low R:FR conditions in the SAM is detected. In the P1, a handful (13) of genes are upregulated in constant shade conditions, including *GA2* (*GA REQUIRING2*), a gibberellin biosynthetic gene homolog and the KNOX gene *LeT6* (sufficient to produce leaflets), suggesting that only a minimal shade avoidance response sustains growth differences between light conditions (**Dataset S6**).

Analysis of genes differentially expressed between the SAM and P1 under constant light conditions (i.e., differential expression by organ type, not light treatment) reveals the same dichotomy between regulators of indeterminacy and differentiation acutely up regulated in the shade P1 after 28 hours (Fig. 6B; Dataset S7). *WUS*, *CLV1*, *LeT6*, three Class I Knox family members, and *PLETHORA* (*PLT*) are among genes up regulated in the SAM relative to the P1 that are significantly enriched for transcription, cell division, and epigenetic-related GO terms (Fig. 6B; Dataset S8). *LYR*/*JAG*, SAWs, and TCPs, as well as photosynthetic GO terms, define genes with increased P1 expression (Fig. 6B; Dataset S9). The association of known developmental regulators of indeterminacy and differentiation with SAM and P1 samples, respectively, attests to the accuracy of our laser microdissection. It also demonstrates the significant overlap between genetic programs associated with the promotion of indeterminacy in the shoot apical meristem with P1 samples acutely exposed to shade for 28 hours, consistent with the increases in leaf serration and complexity we observe during the shade avoidance response (Figs. 1-2; Table 3).

### Shade avoidance and heteroblasty increase leaf complexity through independent pathways

Conspicuously absent from the acute (or constant) transcriptional shade avoidance response we describe in the tomato SAM and P1 are “canonical” shade avoidance response genes, such as *ATHB-2* (*ARABIDOPSIS THALIANA HOMEOBOX PROTEIN2*) (**Datasets S3-S4**) (Carabelli et al., 1993; Devlin et al., 2003). Most of these genes are expressed at sufficient levels that a differential expression call is possible.

Consistent with previous studies, we do observe increased expression of *ATHB-2* in shade in the vegetative apex by qRT-PCR (which includes leaf primordia up to P5-7 and mm in length, as opposed to the SAM and P1 captured by laser microdissection), suggesting that indeed tomato exhibits a canonical shade avoidance response that is restricted to mature organs (Fig. 6C; Steindler et al., 1999). Of course, the most tantalizing hypothesis is that, in mature organs, the canonical molecular shade avoidance pathway cell-autonomously responds to changes in light quality. Yet unknown non-cell autonomous signals would then be transmitted to the shoot apical meristem, where developmental changes in initiating organs take place (Lake et al., 2001; Thomas et al., 2004).

The association of classical developmental pathways regulating incipient leaf specification with the shade avoidance response is unexpected, but intriguing because the phenotypic effects of both pathways are well characterized. Across diverse eudicots, increased expression of KNOX genes (including *LeT6*, up regulated in shade P1) and decreased expression of their negative regulators *PHAN*, SAWs, and *LYR*/*JAG* leads to increased leaf complexity and/or serrations (Janssen et al., 1998; Kim et al., 2003; Kimura et al., 2008; David-Schwartz et al., 2009). A prediction from our data is that shade leaves would be more complex. Indeed, this is the case in domesticated tomato (Fig. 2; Table 3).

Shade is not the only modulator of leaf complexity and the degree of serration, though. Tomato leaves emerging from successive nodes are increasingly complex and serrated (Fig. 2, Table 3) (Chitwood et al., 2012a; 2012b; 2014). Such graded morphological change reflects the temporal status of the vegetative meristem as it transitions from vegetative, to adult, to reproductive fates (Ashby, 1948; Diggle, 2002). Recent work in the Brassicaceae revealed that heteroblastic increases in leaf serration are mediated through temporal licensing by small RNAs of CUCs (CUP-SHAPED COTYLEDONS) through protein interactions with TCPs (TEOSINTE BRANCHED1, cycloidea and PCF transcription factors) (Rubio-Somoza et al., 2014). In tomato, the *CUC2* homolog *Goblet* is required for proper patterning of serrations and leaflet formation (Berger et al., 2009), and the TCP family member *Lanceolate* regulates differentiation of leaf margins (Ori et al., 2007). In response to acute shade, only *Goblet* shows increased expression in the P1, and CUC and TCP family members are not as represented as KNOX genes and other regulators of indeterminacy (**Datasets 3-4**). Despite both shade and heteroblasty mediating increases in leaf complexity and serration, do independent molecular pathways modulate their effects on leaf morphology? This is an especially important question considering the once prevailing hypothesis that changes in shade leaf morphology manifested via alterations in the heteroblastic progression, through prolonged juvenility (Goebel, 1908; Allsopp, 1954; Jones, 1995).

To determine if increases in leaf complexity in shade and the heteroblastic series are modulated by independent pathways at the gene expression level, we performed RNA-Seq analysis of hand-dissected 1) SAMs (consisting of the SAM proper plus 4 leaf primordia) and 2) the P5 at sequential times over a period of 17 days after planting, sampling successive leaf primordia at the same stage of development over the heteroblastic series (Fig. 7A). We note that the hand-dissected P5 used to measure heteroblastic effects does not correspond with the P1 laser microdissected for analysis of shade avoidance, and the subsequent results should be interpreted with this in mind. However, this misses the point: whereas shade avoidance is measured for a consistent developmental stage (P1), measurements of heteroblasty compare successively emerging leaves (that is, different days after planting, dap) at the same developmental time point (P5).

**Figure 7:**
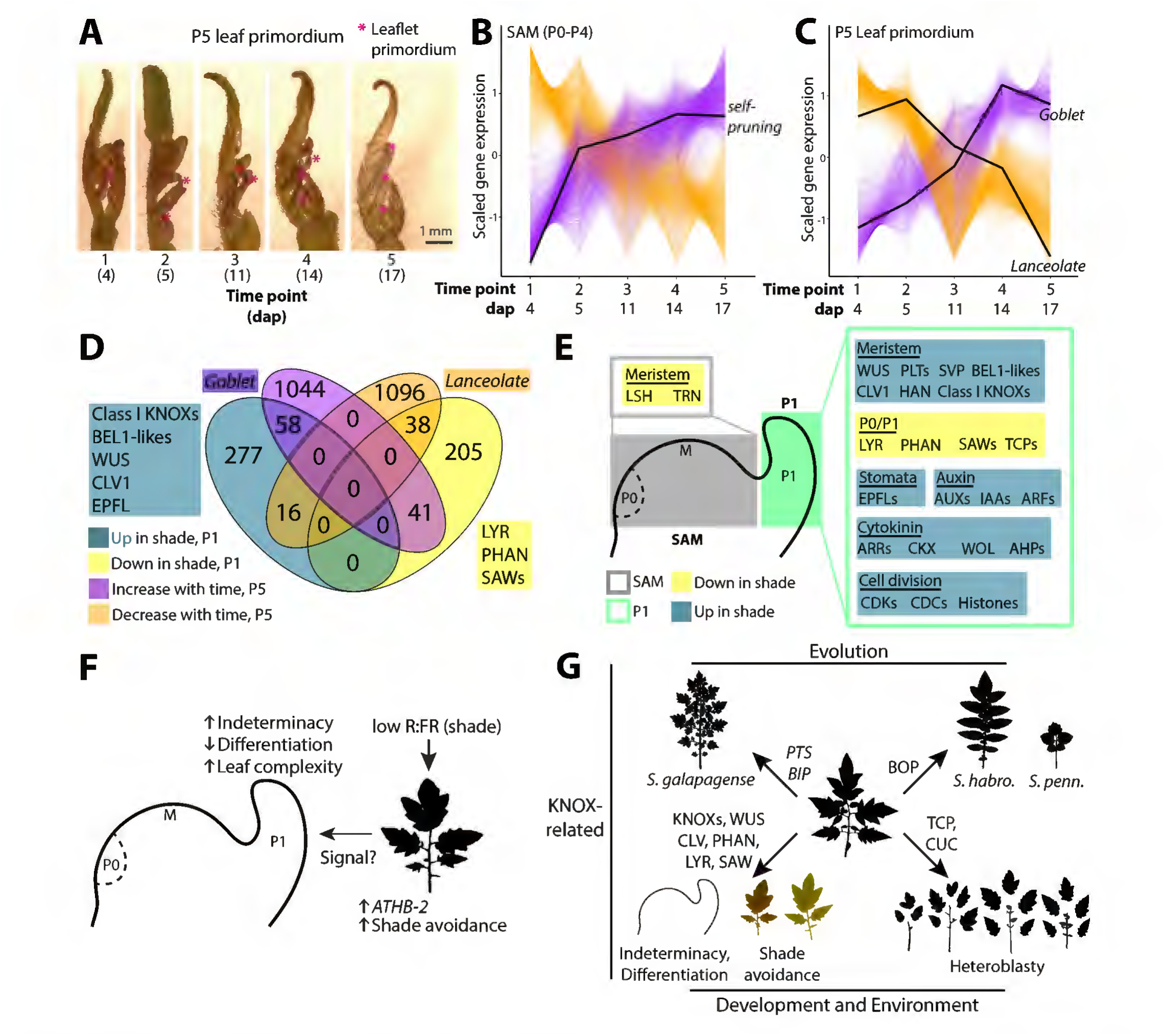
Shade avoidance and heteroblasty both modulate leaf complexity but are transcriptomically distinct processes. **A)** Representative images of P5 leaf primordia that were dissected and analyzed for gene expression using RNA-Seq. At five time points at the indicated days after planting (dap), equivalently staged leaf primordia (P5) and SAM (P0-P4) were dissected. Sampling leaf primordia at the same stage of development separates heteroblastic changes in gene expression (i.e., successively emerging leaves) from the development of individual leaves. Note the increases in leaf complexity of successive leaves, indicated by magenta asterisk. **B-C)** Scaled expression patterns of genes significantly positively (magenta) or negatively (orange) correlated with the heteroblastic series in the **B)** SAM (meristem, plus P0-P4) and **C)** P5. Shown in black are the expression patterns of *self-pruning* in the SAM and *Goblet* and *Lanceolate* in the P5. **D)** Venn diagram of genes that are up-(dark blue) or down-regulated (yellow) upon acute shade exposure in the P1 and genes that are positively (magenta) or negatively (orange) correlated with heteroblasty in the P5. **E)** Sets of genes up regulated upon acute shade treatment (blue) or down regulated (yellow) in the SAM (grey) and P1 (green). **F)** Model for shade-induced developmental change in the SAM. Low R:FR activates the “canonical” shade avoidance response in mature organs, through increased expression of upstream regulators such as *ATHB-2*. A hypothetical signal from mature leaves would increase expression of positive regulators of indeterminacy and effect patterning differences in the P1 (such as increased leaf complexity and serration). **G)** Shade is only one of many factors modulating leaf complexity, including heteroblasty (modulated by CUCs and TCPs) and evolution (modulated by BOP and KNOX-like and BEL-like genes). Shade-induced leaf complexity most closely associates with indeterminacy factors regulating meristem maintenance and leaf differentiation.

Those genes with expression significantly correlated with the heteroblastic series were analyzed further (Fig. 7B-C). In the SAM, we detect 2,171 genes with significantly increasing and 1,797 genes with decreasing expression over the heteroblastic series (**Dataset S10**). *Self-pruning*, a TERMINAL FLOWER1 (TFL1)/FLOWERING LOCUS T (FT) homolog, shows increasing expression over the heteroblastic series (Fig. 7B). Significantly enriched GO terms among genes increasing with the heteroblastic series include small RNA regulation, transcription factor, and photosynthetic terms (**Dataset S11**), consistent with the transition to reproductive development, whereas translation dominates genes with decreasing expression over the heteroblastic series (**Dataset S12**).

In the P5, we detect 1,142 genes with significantly increasing and 1,959 genes with decreasing expression in successive leaves of the same developmental stage (**Dataset S13**). Increasing *Goblet* and decreasing *Lanceolate* expression is consistent with increasing leaf complexity seen in successive leaves (Fig. 7C) (Ori et al., 2007; Berger et al., 2009). Like the SAM, in the P5 genes with significantly increasing expression over the heteroblastic series include GO terms related to small RNA regulation and transcription factor activity (**Dataset S14**). Decreasing gene expression in the P5 is associated overwhelmingly with photosynthetic activities (**Dataset S15**).

Because only 54 genes are differentially expressed in the SAM upon acute shade exposure (**Dataset S3**), we instead focused our efforts analyzing the overlap between differential expression in the P1 upon exposure to acute shade (**Dataset S4**) and changes in P5 gene expression over the heteroblastic series (**Dataset S13**). A majority of genes differentially expressed in acute shade (75.9%) and across the heteroblastic series (93.3%) are not shared between experiments (Fig. 7D). *Goblet* increases in expression upon both acute shade and across the heteroblastic series (consistent with increased leaf complexity for both) but *Lanceolate* is only significantly associated with heteroblasty. Most importantly, significantly increased expression of KNOX family members, *WUS*, *CLV1*, and *EPFL* homologs and decreased expression of *PHAN*/*AS1*, *SAW*, and *Lyrate* is restricted exclusively to the acute shade condition rather than heteroblasty (Fig. 7D).

We conclude that the increases in leaf complexity observed in shade avoiding tomato plants and across the heteroblastic series (Fig. 7B-C) are largely molecularly independent, KNOX and other indeterminacy regulators mediating the former and possibly CUC and TCP homologs mediating the latter. Such a result is consistent with the acute responses to shade observed both morphologically (Fig. 4) and in gene expression (Fig. 5), which argue against heterochronic regulation of development. Further, our results are consistent with classic morphological studies in *Cucurbita* that demonstrated shade does not modulate the heteroblastic progression of leaf forms (Jones, 1995), contrary to prevailing hypotheses of the time (Goebel, 1908; Allsopp, 1954). Although both heteroblasty and shade avoidance affect leaf complexity and serrations (Fig. 2; Table 3), they do so through different associated changes in gene expression.

## Discussion

The shade avoidance response, which allows plants to competitively intercept light and more efficiently use resources when faced with crowding, is inherently morphological in its nature, modifying internode, petiole, and leaf lengths, branching and tillering, the timing of the reproductive transition, and root architecture (Smith and Whitelam, 1997; Steindler et al., 1999; Casal, 2012). In tomato, shade increases leaf complexity and serrations (Figs. 1-2, Table 3) in addition to leaf length, width, area, laminar outgrowth, and stomatal patterning (Figs. 3-4). It is important to distinguish the developmental origins of such plasticity. Do shade induced morphological changes occur beginning with the inception of a leaf, or later? Are the morphological changes heterochronic in nature, modulated through the timing of the heteroblastic series, are do they alter leaf morphology independently? Answering such questions is necessary if a developmental understanding of the plasticity induced by the shade avoidance response is to be achieved.

In tomato, simulated foliar shade alters shoot apical meristem architecture, demonstrating that shade affects the incipient leaf (Fig. 4F). Underlying change in shoot apical meristem size is a strong, transcriptional response to acute shade exposure, especially in the P1 leaf primordium (Fig. 5). Rather than the canonical shade avoidance pathway involved in perceiving and responding to low R:FR light quality in mature leaves, the transcriptional response in the P1 to shade is largely developmental in nature. Shade increases the expression of genes promoting indeterminacy, cell proliferation, or maintaining stem cell identity (such as KNOX, *WUS*, and *CL*V1 homologs and genes modulating cytokinin response) while down-regulating genes involved in leaf differentiation (such as *PHAN* and TCP homologs) (Fig. 7E). Shade-induced expression of canonically responding genes such as *ATHB-2* is restricted to mature organs (Fig. 6C), implying possible non-cell autonomous signals instructing changes in patterning of leaf primordia (Fig. 7F), such as increased leaf complexity and serrations (Figs. 1-2; Table 3).

For over a century, shade-induced changes in leaf morphology have been speculated to be heteroblastic in nature, the morphological changes in leaves arising from successive nodes influenced by nutrition and the amount of available photosynthate (Goebel, 1908; Allsopp, 1954). This hypothesis has recently been revisited, as sugar has been identified as an endogenous signal modulating vegetative phase transition (Yang et al., 2013; Yu et al., 2013). Yet, classical morphological work in *Cucurbita* morphologically separates shade-and heteroblastic-related morphological changes in leaf primordia (Jones, 1995). Our results support the hypothesis that shade-induced morphological change is not mediated through changes in the heteroblastic series. 1) Both morphologically (Fig. 4) and molecularly (Fig. 5) shade avoidance is ephemeral rather than on-going, the latter expected if changes in temporal development had occurred. 2) Measuring the molecular heteroblastic progression in the shoot apical meristem (Fig. 7B) and P5 (Fig. 7C) it is evident that increases in leaf complexity across the heteroblastic series are mediated by TCP and CUC pathways (Rubio-Somoza et al., 2014) rather than the KNOX family members regulating in shade avoidance (Fig. 7D). These results are similar to recent findings that KNOX activity underlies both shade and water-induced increases in leaf complexity in North American Lake Cress (*Rorippa aquatica*, Brassicaceae; Nakayama et al., 2014). The confusion between similar changes in leaves in response to the heteroblastic series and shade avoidance is purely morphological (Fig. 2), reflecting the contributions of distinct molecular pathways (Fig. 7D) to the same trait (Table 3).

The study of plant morphology—how it arises and its potential functions—touches upon all attributes of plant science: development, physiology, genetics, evolution, systematics, and environmental response (Kaplan, 2001). Tomato leaf complexity is modulated by all of these factors (Fig. 7G). Our results put the shade-induced increases in leaf complexity of tomato leaves in a broader evolutionary, developmental, and environmental perspective. Shade avoidance most closely aligns with genes responsible for differentiating leaf primordia from the meristem and patterning leaf complexity, in particular *LeT6* (Janssen et al., 1998; Kim et al., 2003; David-Schwartz et al., 2009), and is distinct from heteroblastic changes mediated by TCPs and CUCs (Ori et al., 2007; Efroni et al., 2008; Berger et al., 2009; Rubio-Somoza et al., 2014), or evolutionary changes mediated by BLADE-ON-PETIOLE (BOP; Ichihashi et al., 2014) or other KNOX-related pathways (*Petroselinum*, *Bipinnata;* Kimura et al., 2008). The morphology of any tomato leaf is the resultant contributions of separate developmental, genetic, and environmental effects (Chitwood et al., 2014; Chitwood and Topp, 2015), modulated by distinct molecular pathways.

## Materials and Methods

### Phenotypic analysis

Tomato seeds (*Solanum lycopersicum*, cv. M82) were washed in 50% bleach for ∼ 2 minutes, rinsed, and placed onto water-soaked paper towels in Phytatrays (Sigma) in the dark for three days at room temperature. Phytatrays were then placed into high R:FR (sun) conditions as described below for another three days. Seedlings were then transplanted into Sunshine Mix soil (Sun Gro) in 9 x 4 sub-divided trays (11’’ x 22’’ in dimensions).

Half the transplanted seedlings were placed into high R:FR (simulated sun) and the other half into low R:FR (simulated shade) conditions. Temperature was adjusted to 22°C and photoperiod to a 16:8 hour light-dark cycle. Lighting consisted of alternating fluorescent (F48T12CWHO) and far-red (F48T12FRHO, peak emission 750nm, Interlectric, Warren, PA) bulbs. High R:FR wavelength ratios were achieved by blocking far-red irradiance with sleeves whereas all bulbs (both normal fluorescent and far-red) transmitted light in the low R:FR red treatment. Shade cover was placed perpendicularly over bulbs in the low R:FR treatment to adjust overall light intensity to match that of the high R:FR condition.

At 20 days after planting (dap), half of the seedlings from each treatment were swapped into the other condition. Measurements were taken every two days, starting 20 dap, and continuing until 34 dap. 17 individuals were measured for each time point for each of four conditions (constant sun, constant shade, sun-to-shade swap, and shade-to-sun swap). At each time point for each condition, plants were randomly selected for analysis.

For leaf area and red-to-green ratio measurements, leaves were arranged under non-reflective glass. Olympus SP-500 UZ cameras were mounted on copy stands (Adorama, 36’’ Deluxe Copy Stand) and controlled remotely by computer using Cam2Com software (Sabsik). ImageJ (Abramoff et al., 2004) was used to threshold, extract, and measure leaf area from photos. The red-to-green ratio of leaves was measured from the abaxial side. Average R, G, and B values (RGB, a digital color model) of pixels in the leaf were recorded by selecting leaves using an appropriate tolerance value to separate them from the white background in Photoshop (Adobe).

For pavement cell and stomatal measurements, dental impression (Provil Novo Light Standard Fast, Pearson Dental Supplies) was applied to terminal leaflets using an application gun and allowed to dry before archiving. Finger nail polish (Sally Hanson, Double Duty) was applied to impressions, allowed to dry completely, removed from the impression, and floated on microscopic slides with water. Water was removed and the nail polish remained affixed to the slide. Micrographs of samples were taken using a standard compound microscope. For each individual impression, two micrographs were taken to insure representative measures. For each micrograph, four pavement cells were traced using Bamboo Tablets (Wacom) in ImageJ (Abramoff et al., 2004) and the area recorded. For stomata, Bamboo Tablets were used to quickly place dots in ImageJ over the feature of interest, followed by custom macros that would count and record the number of features. Pseudoreplication was averaged.

To measure palisade cell size, terminal leaflet tissue was cleared using a 3:1 solution of ethanol:acetic acid for 4 hours, followed by 70% ethanol for 1 hour, and 95% ethanol solution overnight. The following day, leaflets were mounted between two slides in 95% ethanol and images taken with a standard compound microscope. Two micrographs were taken from each sample and for each, the area of four palisade cells measured. Bamboo tablets were used to trace and measure area in ImageJ (Abramoff et al., 2004). Pseudoreplication was averaged.

ANOVA models were fitted for leaf area, palisade and pavement cell size, and stomatal density using the aov() function in R (R Core Team, 2013). Traits were appropriately transformed so that linear values over time were attained. ANOVA models were fitted for constant sun and sun-to-shade swap comparisons as well as constant shade and shade-to-sun comparisons and significance of time, light, and light*time factors determined. Relative growth rate was calculated over two day time spans using the formula (ln(W_2_) – ln(W_1_))/(T_2_ – T_1_) (Evans, 1972), where W_1_ and W_2_ are trait values at times T_1_ and T_2_.

Morphometric analysis was carried out on leaflets grown under simulated sun and foliar shade treatments similar to those described above. Experimental procedures to obtain leaflet outlines are detailed in previous publications, including Chitwood et al., 2012a and Chitwood et al., 2012b describing leaflets from *S. arcanum*, *S. habrochaites,* and *S. pimpinellifolium* accessions and Chitwood et al., 2014 describing leaflets from *S. lycopersicum* cv. M82, *S. pennellii* LA0716, and introgression lines. Linear Discriminant Analysis (LDA) on previously calculated aspect ratio, roundness, circularity, and solidity values, as well as the first three principal components calculated for elliptical Fourier descriptors (Chitwood et al., 2014) was performed using the lda function from the MASS package in R (Venables and Ripley, 2002). LDA was performed to maximize discrimination of plants from different light treatments as a function of different traits for leaflets found across the leaf series. Scaling values defining the contributions of traits and leaflets to discrimination of light treatments from which a plant originates were calculated from scaled values for traits (using the scale function), such that overall trait magnitude did not influence the derived LDA scaling values. The predict function (stats package) and table function (base package) were used (dependent on MASS) to predict the light treatment a plant originated from based on the calculated linear discriminant space. A Mann-Whitney U test (wilcox.test function, R) was used to detect differences in LD values between light treatments.

### Laser capture microdissection and amplification

Tomato plants were grown as described above in either high or low R:FR conditions. Samples either remained in high and low R:FR conditions for the constant sun and constant shade experiment, or high R:FR plants were swapped into low R:FR conditions 28 hrs before harvest. Vegetative apices were harvested 19 dap and were prepared using standard histological procedures as previously described (Belmonte et al., 2013). Briefly, under RNase-free conditions, apices were fixed in ice cold 3:1 ethanol:acetic acid solution, vacuum infiltrated for 1 hour, and fixed overnight rotating at 4°C. Samples were then rinsed three times with 70% ethanol, dehydrated in a graded ethanol series from 70-100% ethanol, and then infiltrated with a xylene series from a 1:3 xylene:ethanol mixture to pure xylene. Samples were incubated with paraffin chips in xylene before numerous paraffin exchanges over 3 days at 60°C. Samples were embedded in disposable rings and blocks and stored for no more than two weeks at 4°C before sectioning. Apices were transverse sectioned at 10μm, one apex per a previously RNase-treated polyethylene napthalate (PEN) membrane slide, and left to dry overnight at room temperature. Slides were subsequently deparaffinized twice in xylene, 1 minute each time.

Approximately 4 shoot apical meristems (SAMs) were microdissected per biological replicate. ∼100,000μm^2^ of 10μm-thick sections were captured for both “SAM” (containing the dome of the SAM and the incipient leaf, or P0) and “P1” (the first separable leaf primodrium) samples. All tissue was collected from sections proceeding (in an apical to basal direction) from the first appearance of the SAM or P1 until their junction. SAM and P1 tissue was collected from the same vegetative apices, such that most SAM replicates have a sister P1 replicate, derived from the same apices. Microdissected tissue was harvested into RNA extraction buffer (RNAqueous-Micro; Ambion) and stored at-80°C until further use.

RNA was isolated following the manufacturer’s instructions, with an additional treatment of the samples on the RNA purification column with RNase-free DNase (1:4 dilution of DNase I in RDD buffer; Qiagen). RNA levels were quantified (Quant-iT RiboGreen RNA Assay Kit; Invitrogen) using a ND-3330 Fluorospectrometer (Nano-Drop). Total RNA was analyzed by microcapillary electrophoresis (RNA 6000 Pico Chip, Agilent 2100 BioAnalyzer; Agilent Technologies) before linear amplification (Ovation Pico WTA System, NuGEN Technologies).

### Laser capture microdissection gene expression analysis

Library preparation for RNA-Seq was performed as in our earlier publication (Kumar et al., 2012) except for the following changes: For second strand synthesis, we added 10μl of cDNA (100-250ng), 0.5μl of random primers and 0.5μl of dNTP. Sample was mixed and heated at 80°C for 2 min, 60°C for 10 sec, 50°C for 10 sec, 40°C for 10sec, 30°C for 10 sec and 4°C for at least 2-5 min. We then added the following (on ice): 5μl of 10 X DNA polymerase I buffer, 31.5μl of H2O and 2.5 μl of DNA Polymerase I, followed by incubation at 16 °C for 2.5 hours. From this point onward the published RNA-Seq protocol (Kumar et al., 2012) was followed, except for the fragmentation time, which in this case was 20 min.

EdgeR (Robinson et al. 2010) was used for differential gene expression analysis. Transcripts for which there were ≤ 2 cpm in < 4 replicates were not considered further. General linearized models were first used to estimate light and tissue effects under constant sun and constant shade conditions (McCarthy et al., 2012), but light was not found to be a significant factor. Therefore, pairwise comparisons between light treatments for a given tissue were made using an exact test based on conditional maximum likelihood methods. For consistency, pairwise methods were used to call differential expression in the constant sun, 28 hr shade experiments as well. FASTQ and processed gene expression data are available from http://www-plb.ucdavis.edu/Labs/sinha/TomatoGenome/Resources.htm

Principal component analysis (PCA) was performed on all replicates using any gene with differential expression between tissues or light in either the constant sun/constant shade or constant sun/28 hr shade experiments using the prcomp() function and visualized using ggplot2 (Wickham, 2009) in R (R Core Team, 2013). Hierarchical clustering was performed on scaled expression values of genes differentially expressed by tissue or light treatment. A distance matrix using the dist() function was calculated followed by hierarchical clustering using the “Ward” method with the hclust() function. Results were visualized using the geom_tile() function from ggplot2 (Wickham, 2009). Promoter enrichment analysis was performed using custom scripts in R with motifs represented in the AGRIS AtTFDB (http://arabidopsis.med.ohio-state.edu/AtTFDB/) (Davuluri et al., 2003; Palaniswamy et al., 2006). Analysis was performed on 1000bp upstream of the ATG translation start site. Abundance of promoter elements in differentially expressed genes was compared to motif abundance in promoters of all considered genes using a Fisher’s exact test allowing for a single mismatch using the Biostrings package (Pages et al., 2013). For select developmental regulators that were 1) differentially expressed and 2) possessed significantly enriched motifs in their promoter, a network was visualized using Gephi (Bastian et al., 2009).

### Heteroblasty gene expression analysis

Tomato plants were grown as described above in a growth chamber at 22°C with 70% relative humidity and a day length of 16 h. P5 leaf primordia and SAMs (consisting of the SAM proper plus 4 leaf primordia) were dissected carefully using razor blade and harvested into RNase-free tubes cooled with liquid nitrogen. This sampling was performed for the following time points: time point 1 (4 dap), 2 (5 dap), 3 (11 dap), 4 (14 dap), and 5 (17 dap). Our experimental design samples successive leaves at the same developmental stage (P5 leaf primordia), measuring effects of the heteroblastic series but not leaf ontogeny.

Heteroblasty RNA-Seq was performed using BrAD-seq, a strand-specific digital gene expression method (Townsley et al., 2015). As brief as possible, tissues were processed and lysed as described in Kumar et al., 2012 with the following modifications: A) Wash volumes of WBA, WBB and LSB were 300 μl each and buffers were chilled on ice prior to use and B) mRNA elution was done into 16 μl of 10 mM TrisHCl pH 8 containing 1 mM beta-mercaptoethanol. Instead of cDNA, mRNA was fragmented using magnesium ions at elevated temperature, and priming of the cDNA synthesis reaction was carried out with 3-prime adaptor polyT priming oligo in presence of 5X Thermo Scientific RT buffer. cDNA synthesis was carried out and subsequently was cleaned and size-selected using Ampure XP beads. After 5 minutes samples were placed on a magnetic tray, supernatant was removed, and pellets were washed twice with 80% ethanol. Residual ethanol was removed and samples were allowed to air dry. Dry pellets were rehydrated by addition of 4 μl of 10 μM 5-prime breath capture adapter. A 6 μl volume of reaction mix containing *E. coli* DNA polymerase I was added to the hydrated pellet, mixed by pipetting and the pre-enrichment library synthesis reaction was allowed to proceed at room temperature for 15 minutes.

The pre-enrichment libraries were washed and size selected using AmpureXP beads present from the previous step and allowed to stand prior to placing on the magnetic tray. Supernatant was removed and pellets were washed twice with 80% ethanol. Pellet was resuspended and placed on magnetic tray. Supernatant was transferred without beads to fresh strip tubes and stored at-20 prior to enrichment by PCR. Samples were amplified in a thermocycler using the program: (98 C 30 seconds, (98 C 10 seconds, 65 C 30 seconds, 72 C 30 seconds) 11 cycles, 72 C 5 min, 10 C hold). Each library sample was run on a 1% agarose gel for size and quantity reference. The remaining 8 μl of enriched library sample was cleaned and size selected using AmpureXP beads and washing twice with 80% ethanol, similar to previous wash steps. The libraries were eluted from the pellet with 10mM Tris pH 8.0 and quantified and pooled as descried in Kumar et al., 2012. 50 bp single end sequencing was carried out at the Vincent J. Coates Genomic sequencing Facility at UC Berkeley.

Gene expression analysis was carried out as described above for the laser capture microdissection dataset to arrive at normalized, log_2_(x+1)-transformed read count values. Replicates were assigned values of 1-5 corresponding to the time point at which they were sampled and the Pearson correlation coefficient and p-value calculated. p-values were multiple test adjusted to control false discovery rate (FDR) using the Benjamini-Hochberg (BH) method. GO enrichment terms for genes up or down regulated over the heteroblastic series in the SAM (P0-P4) and P5 were determined at a 0.05 FDR cut-off value using the “goseq” package in Bioconductor (Young et al., 2010). Venn diagrams were created using VennDiagram (Chen and Boutros, 2011). Unless otherwise specified, all analyses were carried out in R (R Core Team, 2013) and graphics visualized using ggplot2 (Wickham, 2009).

### Sequence submission

The quality filtered, barcode-sorted, and trimmed short read dataset was deposited to the NCBI Short Read Archive under Bioproject accession SRP061929. RNAseq reads from constant sun-shade experiment were deposited under accessions SRR2141260, SRR2141262-SRR2141279. Threads from the transient shift shade experiment were deposited under accessions SRR2141280-SRR2141298, SRR2141300, SRR2141302-SRR2141306, SRR2141314-SRR2141316. The reads from heteroblasty experiment were deposited under accessions SRR2141325-SRR2141369.

## Acknowledgements

Dan Chitwood was a Gordon and Betty Moore Foundation Fellow of the Life Sciences Research Foundation. This work was funded through a National Science Foundation grant (IOS-0820854) to Neelima Sinha, Julin Maloof, and Jie Peng. This work used the Vincent J. Coates Genomics Sequencing Laboratory at UC Berkeley, supported by NIH S10 Instrumentation Grants S10RR029668 and S10RR027303. Bioinformatics was carried out using the NSF-funded iPlant Atmosphere cloud service.

## Author Contributions

DHC, JJH, JNM, and NRS designed the research. DHC, RK, AR, JMP, BTT, YI, CCM, and KZ performed research. BTT contributed new experimental tools. DHC, RK, AR, BTT, and CCM analyzed data. DHC wrote the paper with contributions from other authors.

**Supplemental Data**

**Dataset S1: Morphometric data for leaflets from domesticated tomato and wild relatives grown under simulated sun and foliar shade conditions.** A table providing ID information (“id”), dataset (“dataset”; “arcan”, “habro”, and “pimp” indicating “*S. arcanum*”, “*S. habrochaites*”, and “*S. pimpinellifolium*, respectively, from Chitwood et al., 2012a and 2012b; “il”, “m82”, and “penn” indicating “introgression line”, “*S. lycopersicum* cv. M82”, and “*S. pennellii* LA0716”, respectively, from Chitwood et al., 2014), light treatment (“U”, simulated sun; “H”, simulated shade), and trait values (“ar”, aspect ratio; “circ”, circularity; “round”, roundness; “solid”, solidity, and “pc1-3”) for leaflet types measured in this study. Data is re-analyzed in this study from Chitwood et al., 2012a, Chitwood et al., 2012b, and Chitwood et al., 2014 for meta-analysis purposes, and these publications can be referenced to obtain more complete morphometric information.

**Dataset S2: Scaling values for Linear Discriminant Analyses (LDAs) separating leaflet morphology by light treatment.** A table providing which LDA analysis (“analysis”; “m82”, “*S. lycopersicum* cv. M82”; “il”, “introgression line”; “penn”, “*S. pennellii* LA0716”; “arcan”, “*S. arcanum*”; “habro”, “*S. habrochaites*”; “pimp”, “*S. pimpinellifolium*” accessions), which trait (“trait”), leaf (“leaf”), leaflet type (“type”), and the scaling value (“scale”).

**Dataset S3: Differential gene expression in the SAM between constant sun and 28 hr shade treatments.** A table, providing differential gene expression information between light treatments (constant sun and 28 hr shade) in the SAM, as well as gene annotation information derived from Arabidopsis homologs meeting cut-off requirements. Data is sorted by FDR value. Transcript name (itag), logFC (< 0, up in sun; > 0 up in 28 hr shade), log counts per million (logCPM), unadjusted p value (PValue), FDR (multiple test adjustment controlling for false discovery rate), chromosome, beginning gene position (begin), ending gene position (end), human readable description of best Arabidopsis BLAST hit (human.readable.description), TAIR identifier (Subject.ID), description, BLAST score, e value, and identity of the match are provided for each gene. A worksheet with only differentially expressed genes (FDR < 0.05) has been provided as well, named “Only DE.”

**Dataset S4: Differential gene expression in the P1 between constant sun and 28 hr shade treatments.** A table, providing differential gene expression information between light treatments (constant sun and 28 hr shade) in the P1, as well as gene annotation information derived from Arabidopsis homologs meeting cut-off requirements. Data is sorted by FDR value. Transcript name (itag), logFC (< 0, up in sun; > 0 up in 28 hr shade), log counts per million (logCPM), unadjusted p value (PValue), FDR (multiple test adjustment controlling for false discovery rate), chromosome, beginning gene position (begin), ending gene position (end), human readable description of best Arabidopsis BLAST hit (human.readable.description), TAIR identifier (Subject.ID), description, BLAST score, e value, and identity of the match are provided for each gene. A worksheet with only differentially expressed genes (FDR < 0.05) has been provided as well, named “Only DE.”

**Dataset S5: Over-represented Gene Ontology terms for genes up-regulated in the shade P1.** GO terms that are significantly enriched (p < 0.05, controlling for false-discovery rate) for genes that are up regulated in the P1 upon a 28 hr shift to low R:FR conditions.

**Dataset S6: Differential gene expression in the P1 between constant sun and constant shade treatments.** A table, providing differential gene expression information between light treatments (constant sun and constant shade) in the P1, as well as gene annotation information derived from Arabidopsis homologs meeting cut-off requirements. Data is sorted by FDR value. Transcript name (itag), logFC (< 0, up in sun; > 0 up in constant shade), log counts per million (logCPM), unadjusted p value (PValue), FDR (multiple test adjustment controlling for false discovery rate), chromosome, beginning gene position (begin), ending gene position (end), human readable description of best Arabidopsis BLAST hit (human.readable.description), TAIR identifier (Subject.ID), description, BLAST score, e value, and identity of the match are provided for each gene. A worksheet with only differentially expressed genes (FDR < 0.05) has been provided as well, named “Only DE.”

**Dataset S7: Differential gene expression between the SAM and P1.** A table, providing differential gene expression information between tissues (SAM and P1) from the constant sun and constant shade data. Differential expression was analyzed without regard to light treatment, as a generalized linear model found light to be a non-significant term under constant sun and constant shade conditions (data not shown). Gene annotation information derived from Arabidopsis homologs meeting cut-off requirements is provided. Data is sorted by FDR value. Transcript name (itag), logFC (< 0, up in SAM; > 0 up in P1), log counts per million (logCPM), unadjusted p value (PValue), FDR (multiple test adjustment controlling for false discovery rate), meri.pri (PRI = significantly up in P1; MERI = significantly up in SAM; NA = not significant), human readable description of best Arabidopsis BLAST hit (human.readable.description), TAIR identifier (Subject.ID), description, BLAST score, e value, and identity of the match are provided for each gene. A worksheet with only differentially expressed genes (FDR < 0.05) has been provided as well, named “Only DE.”

**Dataset S8: Over-represented Gene Ontology terms for genes up-regulated in the SAM.** GO terms that are significantly enriched (p < 0.05, controlling for false-discovery rate) for genes that are up regulated in the SAM (ignoring light treatment, in the constant sun, constant shade data).

**Dataset S9: Over-represented Gene Ontology terms for genes up-regulated in the P1.** GO terms that are significantly enriched (p < 0.05, controlling for false-discovery rate) for genes that are up regulated in the P1 (ignoring light treatment, in the constant sun, constant shade data).

**Dataset S10: Correlation of gene expression with the heteroblastic series in the SAM.** A table, providing transcript name (itag), Pearson’s correlation coefficient (r), p value (pval), false discovery rate controlled p value (bh), whether a gene is significantly positively (up) or negatively (down) correlated with heteroblastic series or shows no significant correlation (NS), log_2_(x+1) transformed normalized gene expression values for time points and replication, mean transformed gene expression values at each time point, and gene annotation information.

**Dataset S11: Over-represented Gene Ontology terms for genes positively correlated with the heteroblastic series in the SAM.** GO terms that are significantly enriched (p < 0.05, controlling for false-discovery rate) for genes that are positively correlated with the heteroblastic series in the SAM.

**Dataset S12: Over-represented Gene Ontology terms for genes negatively correlated with the heteroblastic series in the SAM.** GO terms that are significantly enriched (p < 0.05, controlling for false-discovery rate) for genes that are negatively correlated with the heteroblastic series in the SAM.

**Dataset S13: Correlation of gene expression with the heteroblastic series in the P5.** A table, providing transcript name (itag), Pearson’s correlation coefficient (r), p value (pval), false discovery rate controlled p value (bh), whether a gene is significantly positively (up) or negatively (down) correlated with the heteroblastic series or shows no significant correlation (NS), log_2_(x+1) transformed normalized gene expression values for time points and replication, mean transformed gene expression values at each time point, and gene annotation information.

**Dataset S14: Over-represented Gene Ontology terms for genes positively correlated with the heteroblastic series in the P5.** GO terms that are significantly enriched (p < 0.05, controlling for false-discovery rate) for genes that are positively correlated with the heteroblastic series in the P5.

**Dataset S15: Over-represented Gene Ontology terms for genes negatively correlated with the heteroblastic series in the P5.** GO terms that are significantly enriched (p < 0.05, controlling for false-discovery rate) for genes that are negatively correlated with the heteroblastic series in the P5.

